# Large-scale experimental validation of phenotype-guided generative AI for de novo drug discovery

**DOI:** 10.1101/2025.09.13.676062

**Authors:** Jure Fabjan, Joanna M. Wenda, Claire Pecoraro-Mercier, Paula Andrea Marin Zapata, Joerg Wichard, Djork-Arné Clevert, Oscar Méndez-Lucio, David Rouquié

## Abstract

Generative AI is increasingly used to design drugs with desired physicochemical and biological properties. We have previously used the cGAN algorithm trained on cell painting data to generate molecules targeting 10 molecular cancer targets. Here, we provide experimental evidence of the cGAN performance in designing chemical compounds with desired biological activity. We chose a subset of the AI-generated structures for chemical synthesis and tested their activity in a cell painting and a transcriptomic assay. In summary, 88% of the compounds had a pronounced impact on overall cellular morphology, and 37% of the compounds significantly affected transcriptomic pathways associated with the intended target. In a cellular activity assay for an exemplary target, TP53, 6 out of 9 compounds showed potential to modulate TP53 activity. Overall, we show the value of conditioning a generative model on phenotypic readouts for hit discovery of new small molecule drug candidates and pave the way toward AI-guided chemical safety by design.

## INTRODUCTION

Drug discovery is a complex, multistep process, whose length and cost have been steadily increasing in recent years^1–3^. A typical early-stage drug discovery pipeline involves high-throughput screens (HTS) of large small molecule libraries, in which compounds with desired effects are identified as hits, their derivatives re-synthesised, and directed to subsequent screens to optimise their properties^4^. These screens are usually designed with prior biological knowledge on the molecular disease target^4^. Recent advances in computer-aided drug discovery help simplify and accelerate the initial steps of drug screening. Computer algorithms use known drug libraries to model their structure-activity relationships or design a molecule to bind a target based on its known 3D structure. These models allow virtual scoring of large quantities of chemicals for their desired properties and prioritise candidates for synthesis and experimental testing, potentially cutting cost and time on the screening step^5^. Recent advances in machine learning algorithms can bring a further advantage to drug screening by identifying trends and correlations in complex biological data^6^. They can therefore make predictions on how molecular features relate to drug effectiveness, without needing to describe their biochemical interactions explicitly. They are especially effective when trained on datasets containing the outcome of interest^1^.

A specific subtype of the machine learning-based approaches in drug discovery is generative artificial intelligence (AI) – using machine learning models to design chemical structures with desired properties by sampling from learned distributions based on supplied training data. Several generative AI algorithms can be used for creating chemical structures (reviewed in ^7^). Two among them have been especially popular for conditional generation of drug-like molecules: GANs (Generative adversarial networks)^8,9^ and VAE (Variational autoencoders)^10–12^. These models can rely solely on chemical similarity^10,13^, use available information on the target structure^14,15^, or directly use data from biological experiments^8,9,11,12^.

Of the above categories of models, the one using only chemical structure as an input has the narrowest applicability domain (i.e., generation of compounds from their (sub)structure, linker design, etc.). Target-conditioned design aims to create molecules that bind directly to the target of interest by modelling the compound-target interactions. Though this approach was shown to develop good binders, it has two main drawbacks: 1) it needs experimental data of compounds tested in each target of interest, which limits the available training sets, and 2) it does not account for other molecular interactions and the complexity of the biological system, which is critical in assessing the general therapeutic effect. These get better incorporated in the phenotype-conditioned models, which use a multi-dimensional measure of the biological outcome of drug application^7^. The latter case is particularly advantageous since the model extracts patterns from screening data to learn the link between the chemical structure and biological response.

Data from high-dimensional assays allow complex biological responses to be modelled more accurately and the simultaneous analysis of several targets. However, they may also be more costly and difficult to interpret, posing challenges in distinguishing the proper biological response from technical noise and determining their biological relevance. Furthermore, these assays are difficult to standardise on the experimental level, requiring complex normalisation and batch correction strategies. The standard single-target assays are relatively simple, reproducible, and easy to design and validate. However, as they focus on specific interactions without factoring in the complexity of the whole network, using them could result in missing systemic changes and off-target effects. Notably, a suitable target may not be known for some diseases, making a single-target assay difficult to implement^7^.

With an ever-reducing cost of “omic” data acquisition, high-dimensional assays such as transcriptomics, metabolomics, and phenomics make a good choice when training generative AI. They allow the cellular response to treatment to be captured on a system level, providing more holistic information on the compound’s action, possibly including its off-target effects. Furthermore, they are not restricted to a particular target and hence can identify bioactive molecules that function through undescribed mechanisms or affect a few targets coordinatively. Herein, we focus on using imaging data collected in a cell painting assay for the conditioned generation of molecules^9^. This method allows for the assessment of changes in cellular morphology after applying a treatment with a perturbing agent, thanks to the fluorescent staining of cell compartments^16,17^. Of the methods mentioned above, it is the most target-agnostic. It also provides a very favourable cost-per-sample compared to other data modalities, which produce a similar amount of data (e.g., transcriptomics)^18^.

Previously, we trained a conditional generative adversarial network (cGAN) on a set of cell painting morphological profiles elicited by 30,000 small molecules. Through the training, cGAN learned to generate structures of compounds based on the morphological changes they induce in cells^9^. Then, the model was used to create ∼300,000 chemical structures from 31 morphological profiles of overexpression of 10 target genes^9^. In this study, we aim to apply cGAN to a general hit identification scenario. We synthesize 76 compounds from the previously generated set (Table S1) and characterize them using cell painting and transcriptomic GoScreen assay, and for a chosen target (TP53) with a cellular activity assay. To our knowledge, this represents the biggest experimentally validated dataset of AI-generated *de novo* compounds to date^7^. We show that cGAN with subsequent chemoinformatic filters generates predominantly (88%) morphologically active molecules, with a high rate (37%) of on-target-pathway effects. Our results reinforce the idea that using models conditioned on high-dimensional data is a promising strategy for *de novo* design of hit-like molecules.

## RESULTS

### Generated compounds show biological activity

To test the utility of cGAN for hit generation, we selected a subset of the ∼300,000 chemical structures generated in ^9^ for synthesis. First, only chemically plausible structures were selected, followed by the application of filters for desired physicochemical properties and synthesizability. Finally, the top 76 molecules were chosen for synthesis based on their physicochemical properties (Table S1, see Methods for more in-depth description of criteria; note that the number of molecules varies per target) and then tested in biological assays (Figure 1A). In terms of the number of compounds and targets, this is the most extensive study for generating small molecules using ML approaches^7^, including experimental validation.

**Figure 1.**
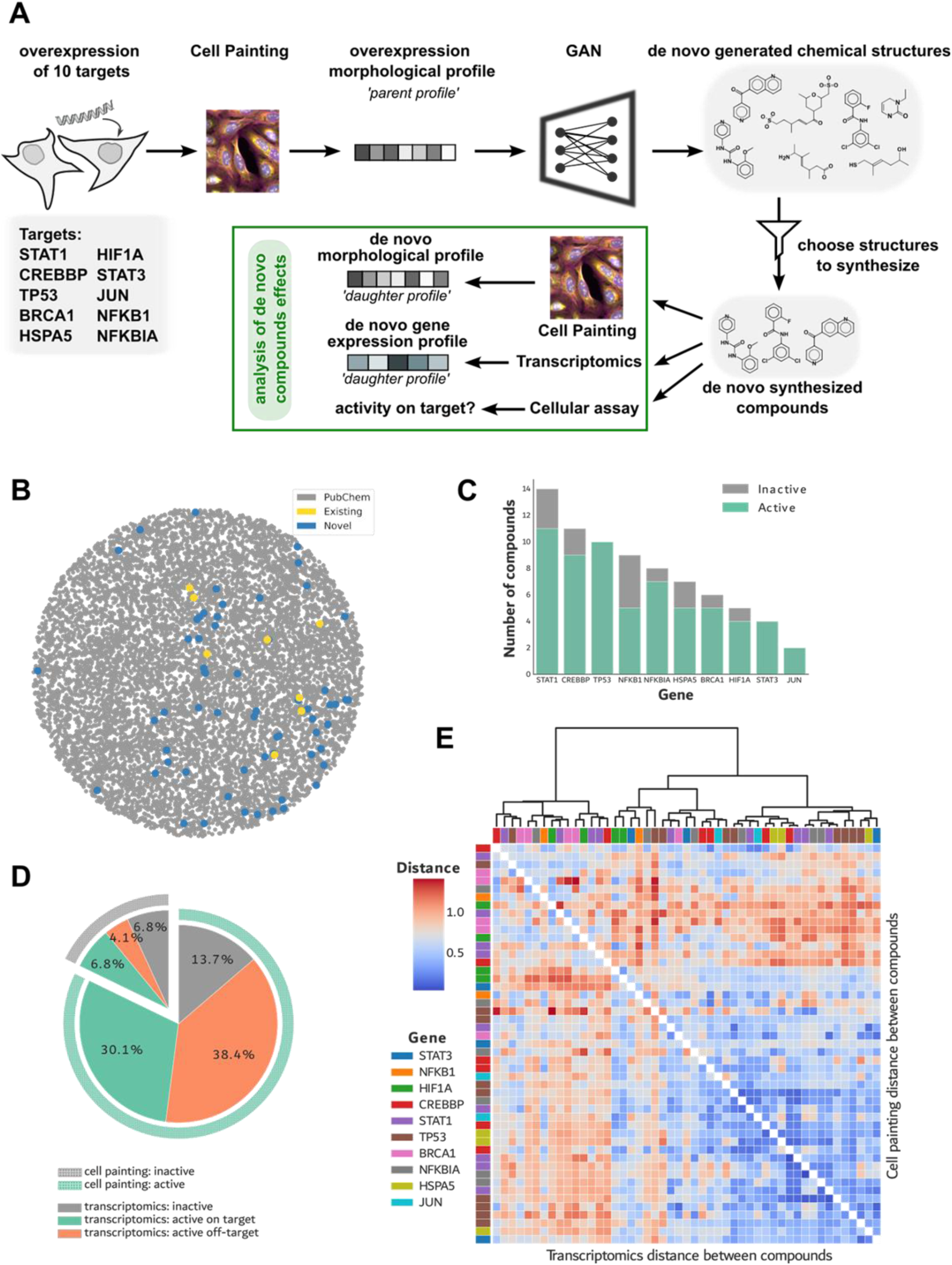
cGAN-generated de novo compounds are chemically diverse and biologically active in chosen validation assays. A) Schematic view of the experiment. Cell painting data from a gene overexpression experiment^19^ was used to generate de novo compounds. A subset of these was selected on chemical and biological constraints (see: Methods) for validation using cell painting, transcriptomics, and bioactivity assays. B) A t-map projection of chemical similarity of compounds relative to the chemical space of compounds available on PubChem. Randomly selected compounds (100,000 molecules) from the PubChem database are depicted in grey. In contrast, de novo compounds are represented in yellow (if the structure matches one previously reported on PubChem) and blue (for structures not previously reported). C) Number of compounds per target classified as active and inactive by their cell painting signature. D) Results of transcriptomic pathway analysis (compounds tested N=73, WikiPathways used for annotation). The inner part of the pie represents the results of the pathway analysis, while the outer ring shows the morphological activity as in panel C. In the outer ring, the morphologically active compounds are represented in green, and the inactive compounds in grey (using the Kendall test on Mahalanobis distances). On the inner circle, compounds perturbing at least one pathway engaging their target gene are shown in green, compounds affecting only pathways unrelated to the target gene are colored orange, and compounds without differentially expressed genes are shown in grey. E) Heatmap of de novo compounds’ similarity measured as the cosine distance between profiles. The upper-right triangle shows the distance between compounds in cell painting data, while the lower-left triangle shows distances in transcriptomic data. Rows and columns are annotated with the gene that was overexpressed to generate the compound. Both rows and columns are ordered based on the transcriptomic similarity (dendrogram on top).

As reported by ^9^, cGAN can generate compounds representing a sampling of diverse chemical space. We first searched for the structures of the 76 synthesized compounds on the PubChem database, revealing that most (48/76) were novel chemical entities. Furthermore, no generated compound was part of the dataset used in cGAN training^9^. Then, we compared the structures of the synthesized molecules to a sample of PubChem molecules as the background reference for the known chemical space (Figure 1B). Even after applying the synthesis selection criteria, the compound sample covers a significant portion of the known chemical space, similarly to observations made by Marin Zapata et al.^9^. The sampling of completely novel compounds is not linked to any specific portion of the chemical space (Figure 1B). These observations indicate that the model correctly learns chemistry rules and can produce novel compounds different from the ones in the training set.

We then assessed the biological activity of the generated compounds. As the model generates molecules from cell painting profiles, we define a bioactive compound as any molecule capable of inducing morphological changes in the cell. We tested the generated compounds at three concentrations in a cell painting assay on U2OS cells (Figure 1A). We defined the activity as a dose-dependent increase in morphological distance from the negative control (Supplementary Figure S2). By applying the Kendall tau test to the Mahalanobis distances (MD), we observed dose-dependent response in 88.16% (67/76) of compounds (Figure 1C and Supplementary Figure S2). The activity was not constrained to a specific target. In fact, all targets had at least two active compounds, and all compounds generated for TP53, STAT3, and JUN were bioactive.

### Generated compounds affect specific pathways

With most of the generated molecules being bioactive, we next characterized their induced biological effects in the U2OS cell line. We define a parent profile as the morphological profile of the genetic perturbation used to condition the design of a specific compound (Figure 1A), assuming that the corresponding *de novo* compounds will induce a similar response in the U2OS cells. In our case, parent profiles are caused by the overexpression of specific pharmacological targets in those cells (see: Methods). We first checked if the generated molecules reproduce the morphological response of their respective parent profile. If this is the case, we anticipate a higher degree of similarity between a compound and its parent profile than a randomly selected parent profile from a different compound. To assess this, we employed a permutation test to evaluate if distances of the compound morphological profiles to their respective parent profile are smaller than distances to unrelated profiles from the overexpression dataset. This test showed that ‘true matches’ are significantly closer to each other than expected by chance (p-value of 0.004), indicating a general similarity of the morphological profiles of the generated compounds to their parent profiles.

As the parent profiles resulted from the overexpression of specific molecular targets, we checked if the morphological similarity between the daughter (i.e., profile from the generated compound) and parent profiles can be attributed to the perturbation of genes of interest. For this, we performed a transcriptomic assay (GoScreen) on U2OS cells treated with the generated compounds at the concentration that produced a maximum morphological change or the lowest concentration at which the effect plateaued. After subjecting the data to quality control (see Methods), 73 compounds were used in further analysis. Only around 2.6% (2/73) of the generated compounds significantly affect the mRNA level of expression of their respective target gene. Such a small rate is unsurprising, as cGAN generates compounds based on gene overexpression-induced morphology. As such, the model is agnostic of which gene is being targeted and can generate compounds that reproduce the morphology acting on downstream targets. Furthermore, the effect can be exhibited through target activation - mimicking functional overexpression without directly affecting the expression level of the target.

As these mechanisms should still produce changes in the pathways in which the target plays a role, we expanded the on-target testing to the level of pathways. We found that 37% (27/73) of the tested compounds perturb at least one pathway containing the target gene. Notably, 20.5% (15/73) of compounds did not induce any significant change in the gene expression levels (Figure 1D). Overall, 93.2% of compounds tested in both assays exhibited an effect either transcriptomically, morphologically, or both. More specifically, 13.7% of compounds affected only morphology, while 10.9% showed activity only in the transcriptomic data (Figure 1D). These discrepancies highlight the importance of using multiple data modalities to verify compound effects.

We hypothesized that the generated compounds conditioned to mimic the morphology induced by the overexpression of a given gene should produce similar biological effects. To gauge the compound similarity, we used two data modalities – cell painting and transcriptomics, to perform hierarchical clustering of compound profiles (Fig. 1E). The clustering indicates a somewhat weak cosine similarity among compounds designed to target the same gene. This dissimilarity could be attributed to the fact that not all generated compounds fully match the condition guiding the generative process, based on the cGAN discriminator score. In fact, only 35% of the generated compounds match the condition to 70% or more. However, some compounds conditioned with the overexpression pattern of the same target – most notably 5 out of 10 TP53-targeting compounds, aggregate within the same cluster and show similarity in terms of induced morphology and transcriptomic profiles. Interestingly, similar distance patterns can be distinguished across both data modalities for most compounds.

### Generated compounds show on-target enrichment

We decided to examine in more detail the generated compounds designed to recapitulate morphological profiles of TP53 overexpression (TP53-generated compounds). The choice was made because of their relatively close clustering on the heatmap, the representative number of compounds for this target (10), and the availability of cellular assays for p53 functions. First, we used the Brunner-Munzel test to see if the compounds’ effects (at phenotypic and transcriptomic levels) show higher similarity within their group compared to their similarity to the rest of the compounds. The morphological profiles of the TP53-generated compounds are significantly more similar (p-value of 0.001) compared to all other generated compounds that showed morphological activity. On the other hand, the average distance between transcriptomic profiles of the TP53-generated compounds is slightly higher than their average morphological distance (Figure 2A). Using the same test to evaluate the transcriptomic profile similarity showed no tendency for the transcriptomic profiles to aggregate (p-value of 0.22).

**Figure 2.**
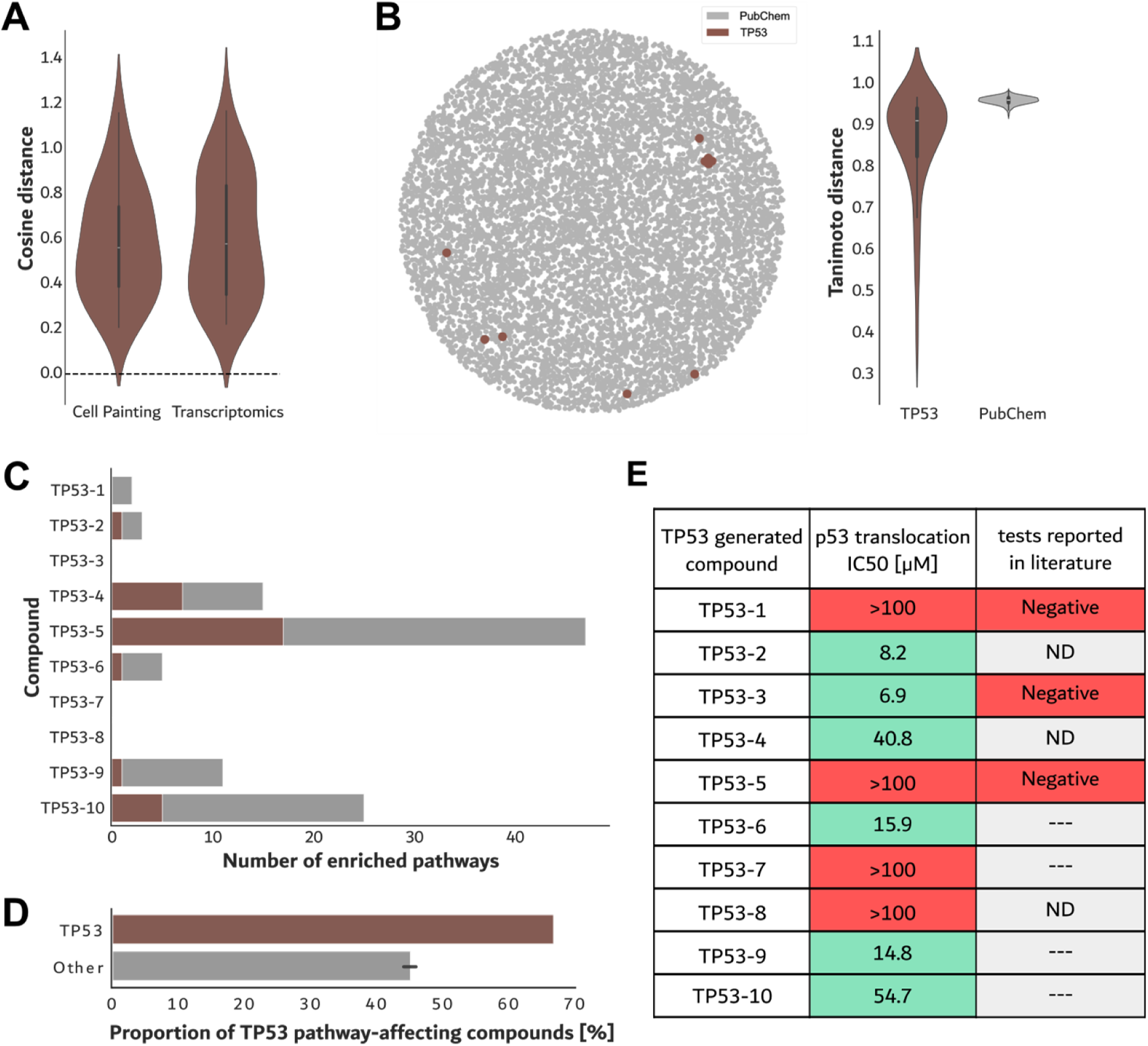
Evaluation of compounds from selected overexpression profiles: TP53 and CREBBP. A) Distribution of pairwise cosine distances between compounds generated from the TP53 overexpression profile. The distance is calculated from cell painting (left violin) and transcriptomics (right violin). B) Left: a representation of chemical similarity of TP53 de novo compounds (in brown) relative to PubChem chemicals (100,000 randomly selected molecules depicted in grey, as in Fig. 1B). Right: pairwise Tanimoto distances between chemical structures in the indicated category. To generate the PubChem distribution, 10 random compounds were repeatedly sampled, and the mean pairwise distance was calculated. C) Number of perturbed pathways (annotation with WikiPathways) in transcriptomic experiment for each TP53 de novo compound. The pathways containing TP53 are depicted in brown, while all other pathways are shown in grey. D) Proportion of compounds that induce a change in at least one pathway related to TP53. For the compounds with a non-TP53 parent profile (‘other’), the bar represents an average of 100 samplings of random 10 compounds. Error bars depict the standard deviation of the mean. E) Table with the results of cellular activity assays for de novo compounds generated from TP53 overexpression. “p53 translocation” refers to the nuclear translocation assay from this study. In contrast “tests reported in literature” refer to reports extracted from PubChem (for details see Table S2), compounds inducing a response are marked in green, and compounds not inducing a measurable effect within the concentration range tested are marked in red (max conc. for the p53 translocation was 100 μM); ND – no data, ‘---’ – compound structure not reported in PubChem.

Next, we looked at the TP53-targeting compounds in terms of their chemical similarity. They are sampled from a diverse but constrained subset of the chemical space (Figure 2B). Comparing the distribution of pairwise Tanimoto distances to the distribution of random samples from PubChem shows that the generated compounds tend to have a comparably high distance and a wider distribution (Figure 2B). This indicates that cGAN tends to explore a relatively wide chemical space even when it generates molecules from a narrow biological profile.

Analysis of transcriptomic pathways revealed that 6 out of 9 (67 %) of the transcriptomically active TP53-generated compounds significantly affected at least one pathway associated with the target gene (Figure 2C). However, none of the compounds modulate the mRNA expression level of TP53 itself. Importantly, only 46% of unrelated compounds (designed to target other genes than TP53) perturbed at least one TP53-associated pathway (Figure 2D). These results show that cGAN-designed compounds are enriched to affect the target of interest.

Finally, we assessed the compound effects on p53 more directly, measuring p53 translocation to the nucleus as an indication of its activation. Six out of ten compounds showed inhibitory effects on nuclear translocation, with IC_50_ values ranging from 6.9 to 54.7 μM (Figure 2E). These findings confirm that these six compounds modulate p53 activity, though we emphasize that the method cannot measure a direct binding to p53. Furthermore, our results suggest that the compounds do not merely affect the p53 pathways by only inducing cellular stress, such as DNA damage or oxidative stress, since these typically lead to p53 nuclear accumulation rather than its inhibition.

While an inhibitory effect might seem counterintuitive given that the compounds were designed based on TP53 overexpression, predicting the directionality of the increased expression impact on p53 subcellular localization is challenging due to the complexities of its regulatory network^20,21^.

Interestingly, the nuclear translocation results are mostly concordant with the transcriptomic analysis, as most compounds positive in the translocation assay (five) also showed changes in the expression of p53-associated pathways. In parallel, we searched the PubChem database and found that six of our generated compounds had already been reported in this public repository. Three compounds (TP53-1, TP53-3, and TP53-5) had been previously tested in high-throughput screens for their ability to bind TP53 modulators but were found inactive. Notably, one of these compounds (TP53-3) affected p53 nuclear translocation without influencing any p53-associated transcriptomic pathways, while a second compound (TP53-5) perturbed multiple p53-associated pathways without affecting its translocation. The discrepancies between the transcriptomics and translocation assays compared to PubChem high-throughput data may be attributed to several factors, including differences in compound concentrations, cell lines used, treatment durations, and assay readouts.

Overall, our in-depth analysis of TP53-generated compounds demonstrates that cGAN can effectively generate compounds that influence the behaviour of a target of interest, highlighting the potential of this framework for hit generation. Nonetheless, these compounds should be evaluated through multiple downstream tests to elucidate further the mechanisms of action by which they modulate their target.

## DISCUSSION

In this work, we validated *de novo* designed molecules generated with cGAN conditioned on cell painting profiles of pharmacological targets overexpression. We focus on several cancer-relevant targets expanding from the usual case of a single target, tested in previous studies of this type. We show that the model, followed by additional chemical and biological filters, can be applied to produce mostly chemically novel and biologically active compounds. Although most generated compounds do not directly influence the expression level of their intended target, they have a high rate (37%) of affecting relevant transcriptomic pathways. For a chosen target (TP53), we demonstrate that these on-target pathway effects are specific - they arise more often for compounds designed to act on TP53 than unrelated compounds.

Furthermore, the general phenotypic effects elicited by TP53-designed compounds are significantly similar. We have shown that conditioning generative models on high-dimensional biological data is a promising strategy for *de novo* design of hit-like molecules, an essential milestone on the road to practical AI-assisted drug design. Multiple further steps still need to be taken before that is achieved. A standard generative benchmark task should be established for the selection of the best-performing model. Furthermore, including additional guiding dimensions such as bioavailability and safety profile is beneficial.

## METHODS

### Choosing chemical structures to synthesise and chemical synthesis

Among the hundreds of thousands of compound structures proposed by the model, we selected a series of chemicals for each conditioned target and applied various filtering criteria. Specifically, Lipinski rules and the predicted potential of the compounds to enter cells were applied, as assessed using an in-house QSAR model trained on Caco-2 cell permeation assay data. The selected 76 de novo-designed compounds were synthesized and subcontracted to Sundia (Shanghai, China). All synthesized compounds were confirmed to have a minimum purity of 90%.

### Cell Painting assay

#### Cell Culture and Cell seeding

All experiments were performed on the human osteosarcoma cells (U2OS) obtained from ATCC (ref.: HTB-96, lot: 70025046). Seeding the microplates was performed directly from a frozen cell stock (assay-ready cells). The cell bank was prepared in house as follows: the vial of U2OS cells obtained from ATCC (ref.: HTB-96, lot: 70025046) was thawed and cultured in the McCoy’s 5A Modified Medium with GlutaMAX™ Supplement (Thermo Fisher, ref: 36600021) supplemented with 10% Fetal Bovine Serum (Gibco, ref.: 16000044) and penicillin/streptomycin mix (Sigma Aldrich, ref: P4458) in standard conditions (humidified incubator, 37°C, 5% CO_2_). The culture was subsequently expanded. The passages were performed with trypsin (Thermo Fisher, ref. 25200056) when the cells reached 80% confluency. The number of cells for re-plating was estimated using trypan blue (Sigma, ref.: T8154) staining and an automatic cell counter (Countess II, Thermo Fisher). At internal passage three (P3), the cells were cryopreserved using complete media with 10% DMSO to obtain a master bank. One vial of the master bank was then used to obtain a working bank by expanding the cells and cryopreserving them at internal passage 6 (P6), which had 4 million cells per vial. The frozen stocks were stored in an ultra-low temperature freezer (-150°C). For seeding the microplates (Greiner BioONE CELLSTAR µCLEAR®; ref: 781091), one vial of the working bank was thawed in the water bath (37°C), the cell suspension was added to 10 ml of pre-heated complete media, and centrifuged (5 min, 120xg). The cell pellet was resuspended in 175 mL of complete medium and placed in a round bottle with a magnetic stirrer. Cell suspension was distributed with Multidrop (Thermo Fisher) into the 36 µl/well microplates, corresponding to around 820 cells/well. The cells were then incubated in an automated incubator (Cytomat 2, Thermo Fisher) for 24 h until treatment. Four experimental replicates were performed, each originating from a separate cell vial (P6). The replicates were processed in two batches, on two different days.

#### Compound treatment

The proprietary test compounds were obtained from internal compound logistics at Bayer Crop Science in powder form in 96-well deep-well plates compatible with the automation setup. All test compounds were dissolved in DMSO (dimethyl sulfoxide) to create 100 mM stock solutions and subsequently diluted in DMSO to create chemical dose plates at the following concentrations: 100 mM, 31.6 mM, and 10 mM. The compound preparation was performed using the Viper liquid handler (Synchron). Control compounds were purchased from the external suppliers and processed manually (dissolved, diluted, and placed in dose plates). The chemical dose plates were prepared separately for each of the four experimental replicates and then preserved at -20°C until use. The compound plates were thawed on the day of the treatment (24h post-cell seeding), and the compounds were added to the cell plates as follows: 1 µl of each compound solution was diluted in 100 µl of complete cell medium (intermediate dilution: 1:100), then 4 µl of this solution were added to each well of the cell plate, containing 36 µl of media (1:10 dilution). The test concentrations in the well correspond to: 10 µM, 31.6 µM, and 100 µM. Final DMSO concentration in cell media was 0.1%. The cell plates were exposed to the test compounds for 48 h.

#### Staining

The fixation and staining were performed with an automated setup. For staining, the PhenoVue JUMP kit was used (Perkin Elmer, ref.: PING23) according to the manufacturer’s instructions and following good practices outlined in ^22^. Mitotracker solution was distributed with Multidrop, 20 µl/well, to the final concentration of 500 nM. The plates were incubated with the dye for 30 minutes at 37°C. Next, 20 µl of 16% PFA (Thermo Fisher, ref.: 28908) were added to each well, and the cell plates were incubated at room temperature (25°C) for 20 minutes. The fixation solution was removed, and the plates were washed 1x with 80 µl of HBSS (Gibco, ref.: 14065-056). Blue Washer (BlueCatBio) was used for washing steps and for the distribution of the staining solution (20 µl/well). The composition of the staining solution was as follows: HBSS, 1% BSA, 0.1% Triton X-100, 43.7 nM PhenoVue Fluor 555 – WGA; 48 nM PhenoVue Fluor 488 - Concanavalin A; 8.25 nM PhenoVue Fluor 568 – Phalloidin; 1.62 µM PhenoVue Hoechst 33342 Nuclear Stain; 6 µM PhenoVue 512 Nucleic Acid Stain. The plates were incubated at room temperature (25°C) for 30 min, and after this time, washed 3x with 80 µl of HBSS, before being sealed with aluminium foil.

#### Image acquisition

The images were recorded with an ImageXpress Micro 4 microscope (Molecular Devices) using a 20x air objective (NA=0.45) and 2x2 camera binning. Nine centrally placed fields of view, corresponding to a total area of 2163 µm x 2163 µm, were recorded per well. For each field of view, images were recorded in five channels, in the following order: Cy5, Texas Red, Cy3, GFP, and DAPI. The Z-offset and exposure times were adjusted for each channel.

### GoScreen (transcriptomic assay)

#### Cell seeding, treatment, and lysis

96-well collagen-coated microplates (Corning 96-well BioCoat, Ref.:356650) were seeded with U2OS cell suspension prepared directly from a thawed vial of a working bank (similarly to the procedure for the Cell Painting assay). The plating density was 6500 cells/well (90 µl of the standard growth medium). The cells were incubated for 24 h prior to treatment.

Internal compound logistics at Bayer Crop Science supplied the desired number of compounds in a powder form in 96-well deep-well plates. The test compounds were dissolved in DMSO to achieve the desired concentration. Each compound was tested at a single concentration in three technical replicates (placed on different chemical plates). For treatment (24 h after cell seeding), compounds were added to the cell plates stepwise, similar to the procedure described for the Cell Painting assay (intermediate dilution in cell media: 1:100, then 1:10 dilution). Final DMSO concentration in treated cell plates was 0.1%. The cell plates were incubated with the test compounds for 48 h at 37°C.

After 48 h, the plates were washed twice with ice-cold PBS (100 µl/well) using a MultiFlo FX device (BioTek). Then, 100 µl of RLT Plus buffer (Qiagen, Ref.: 1053393) was added. The plates were sealed and immediately transferred to -80°C until further processing. They were then sent to Eurofins (Aarhus Denmark) for the GoScreen Human assay (Thermo Fisher Scientific). The GO Screen Human Assay platform is a 384-well DNA microarray plate technology measuring ∼20,000 human genes associated with the Gene Ontology Consortium.

#### p53 translocation assay

The p53 translocation assay was performed at Eurofins with the PathHunter NHR Assays (the cell line used: PathHunter U2OS p53 Nuclear Translocation Cell Line, cat. no. 93-0757C3). Assay performed in dose response using 10 concentrations within the range 0.0051-100 μM.

#### Transcriptomic data processing

The data acquired using the GoScreen platform was analysed using R and its Affy package^23^. The raw files were loaded into a custom script, and the probe intensities were normalized with SST-RMA implemented in ThermoFisher Analysis Power Tools. Next, outliers were manually identified based on the distribution of signal intensities and their position on a PCA plot. This quality control process removed six samples in total. The retained samples were re-normalized, and the control and low-intensity probes were removed.

#### Transcriptomic data analysis

Limma package^24^ was used to fit a linear model comparing the treatment to the DMSO, with the plate as a covariate. For each treatment, the differentially expressed genes were identified as those with an adjusted p value equal to or lower than 0.05 and an absolute fold change higher than 1.5. The differentially expressed genes were queried for the target genes of each compound.

#### Pathway analysis

We performed the pathway analysis on the transcriptomic results using gseWP from the clusterProfiler package. The function performs GSEA analysis on WikiPathways^25^. The results were extracted for all measured genes, including their fold change, for every compound, and used in the analysis. After the analysis, we checked for every compound to see if the target gene was present in any of the affected pathways. To perform this comparison, we extracted the genes involved in each pathway and searched for the gene of interest. As some compounds did not perturb any gene, the procedure resulted in three classes: 1) compounds affecting at least one pathway containing the target gene, 2) compounds eliciting differential expression of at least one gene, but not affecting any pathways containing the target gene, and 3) compounds without any differentially expressed gene. Furthermore, the presence of TP53 was searched in the affected pathways for all compounds. Finally, the result tables were exported into an Excel file to be imported into the Python environment for further processing/plotting.

#### Cell painting feature extraction

The images were segmented, and the morphological features extracted as described by ^26^. For the comparison of the data to the overexpression dataset (cpg0012-wawer-bioactivecompoundprofiling), the latter was fetched from the JUMP (Joint Undertaking for Morphological Profiling) Cell Painting Gallery^19,27^. To relate the datasets better, the images from both were processed using the same pipeline. After obtaining the per-well features for both experiments, they were loaded into the Python environment using Jupyter Notebook. Next, a single column containing invalid values and all columns (features) that were not previously used in cGAN training^9^ were removed.

#### Cell painting data processing

During initial optimization of cell painting experiments (not shown), the wells on the edge of the plate yielded unreliable and variable data (profiles). Therefore, negative control treatment (DMSO) was always applied to the wells in the plate’s first and last rows and columns. DMSO was further applied to either 67 or 70 wells (depending on the plate arrangement). Acquired images from the wells on the plate edge were used in the illumination correction step [see ^26^ for details], but extracted features were excluded from further analysis. Next, we removed samples that deviated too much from other replicates based on the cell count normalized to the plate DMSO. Furthermore, we manually identified and removed samples based on their deviation from other replicates on a PCA plot. The described process removed four samples (compound at a specific dose) out of 228, all DMSO samples on the edge of the plates, and an additional four from other positions, resulting in 544 negative control samples used in the analysis. The remaining data was normalized to the plate DMSO using robust z-scores. Subsequently, a feature selection process was performed on the normalized data using PyCytominer^28^. More specifically, blocklisted features were removed, and correlation and variance thresholds and noise removal were used. Moreover, all features measuring the correlation between different channels (column names starting with “Correlation”) were removed. This procedure retained 270 features.

#### Morphologically active treatments

The Mahalanobis distance to the DMSO on the same plate was calculated using the SciKit-Learn package^29^ for each treatment. The distance values were fed into the Kendall tau test implemented in the SciPy package^30^. We tested the alternative hypothesis of greater and classified compounds as positive if their p-value was strictly lower than 0.1. The p-value threshold was selected based on the visual inspection of the distance dose-response graphs (Supp. Fig. 1) to ensure that the threshold was appropriate to separate the compounds with and without apparent dose-response effect.

#### Comparison to the parent profiles

Using a permutation test, we tested whether the distance between the morphological profiles of compounds and their parent profiles could have arisen by chance. For this purpose, we related a mean morphological profile of a single compound concentration to its parent profile. We selected the concentration for every compound at which the highest deviation from the negative control was elicited using the Mahalanobis distance. If multiple concentrations had the highest deviation, we took the lowest of these concentrations. We first calculated the sum of cosine distances between compound and parent profiles. Next, we performed 10,000 permutations by scrambling the pairings between compounds and parent profiles and calculating the sum of distances for every permutation. The p-value was then calculated as the proportion of the permutations with the same or lower sum of distances compared to the correct pairing of profiles.

#### Within-target similarity

The Brunner-Munzel test was used to check whether the compound morphological profiles from the same group are more similar to each other than to the compounds generated on a different target. The test implementation was taken from ^31^.

#### Merging morphology and transcriptomic results

The results from the transcriptomic analysis were imported into Python. Next, genes were filtered to retain only the ones where at least two compounds had a p-value equal to or lower than 0.01 and an absolute log2 fold change equal to or higher than 2. The application of the filter resulted in a list of 783 retained genes. As the transcriptomic assay was performed using a single concentration per compound, we matched the concentrations on the cell painting. Next, the cosine distance matrix between compounds was calculated for each data modality, and the lower-left transcriptomic and upper-right cell painting matrices were joined. Only the transcriptomic matrix was used to calculate the linkage for the dendrogram shown at the top of the heatmap in Figure 1E. Finally, the correlation between the results obtained in both modalities was calculated by taking the upper-right triangle from the two distance matrices, flattening them, and calculating the Spearman correlation between them.

#### Chemical space

For the background chemical space, a random 100,000 organic molecules were selected. SMILES from PubChem and generated molecules were imported into Python and converted into a 1024-dimensional MinHash fingerprint (MHFP) using the Mhfp package^32^. Tmap^33^ was used to create a low-dimensional representation of the merged set of fingerprints. The plotting was performed using the Seaborn package^34^.

#### Tanimoto distance

The Tanimoto distance was calculated from the MHFP fingerprints using the Mhfp package^32^. The pairwise distance between all TP53-targeting molecules and all CREBBP-targeting molecules was also calculated. For PubChem, we selected a random 10 entries, the same as TP53-targeting molecules, calculated the pairwise distances, and averaged them. This sampling was repeated 1000 times to produce a reference distance distribution used for PubChem.

#### General processing and analysis

In the analysis of the transcriptomic data, we used R v4.2.3^35^ with Affy v 1.76.0^23^ to process the raw data, Limma v 3.54.2^24^ to analyse the data, and biomaRt v 2.54.1^36^ and clusterProfiler v 4.6.2^37^ for pathway analysis. For cell painting and integration of both data modalities and all the plotting, we used Python v3.10.13. For data handling, we used Numpy v1.26.4^38^ and Pandas v2.2.0^39^, while for already implemented processing and analysis calculations, we used PyCytominer v0.2.0^28^, SciKit-Learn v1.4.1^29^, and SciPy v1.12.0^30^. We used RDKit v2024.03.1^40^, Mhfp v1.9.6^32^, and Tmap v1.0.6^33^ to handle chemoinformatic data. For plotting, we used Matplotlib v3.7.2^41^ coupled with Seaborn v0.13.2^34^.

## ACKNOWLEDGEMENTS

The authors thank Fabrice Camilleri and Xabier Cendoya Garmendia for valuable discussion and suggestions on Cell Painting data analysis, Thomas Auda and Fabien Labbal for technical assistance, and Ariane Bassignani for advice on GoScreen.

## Supplementary Information

**Figure S1:**
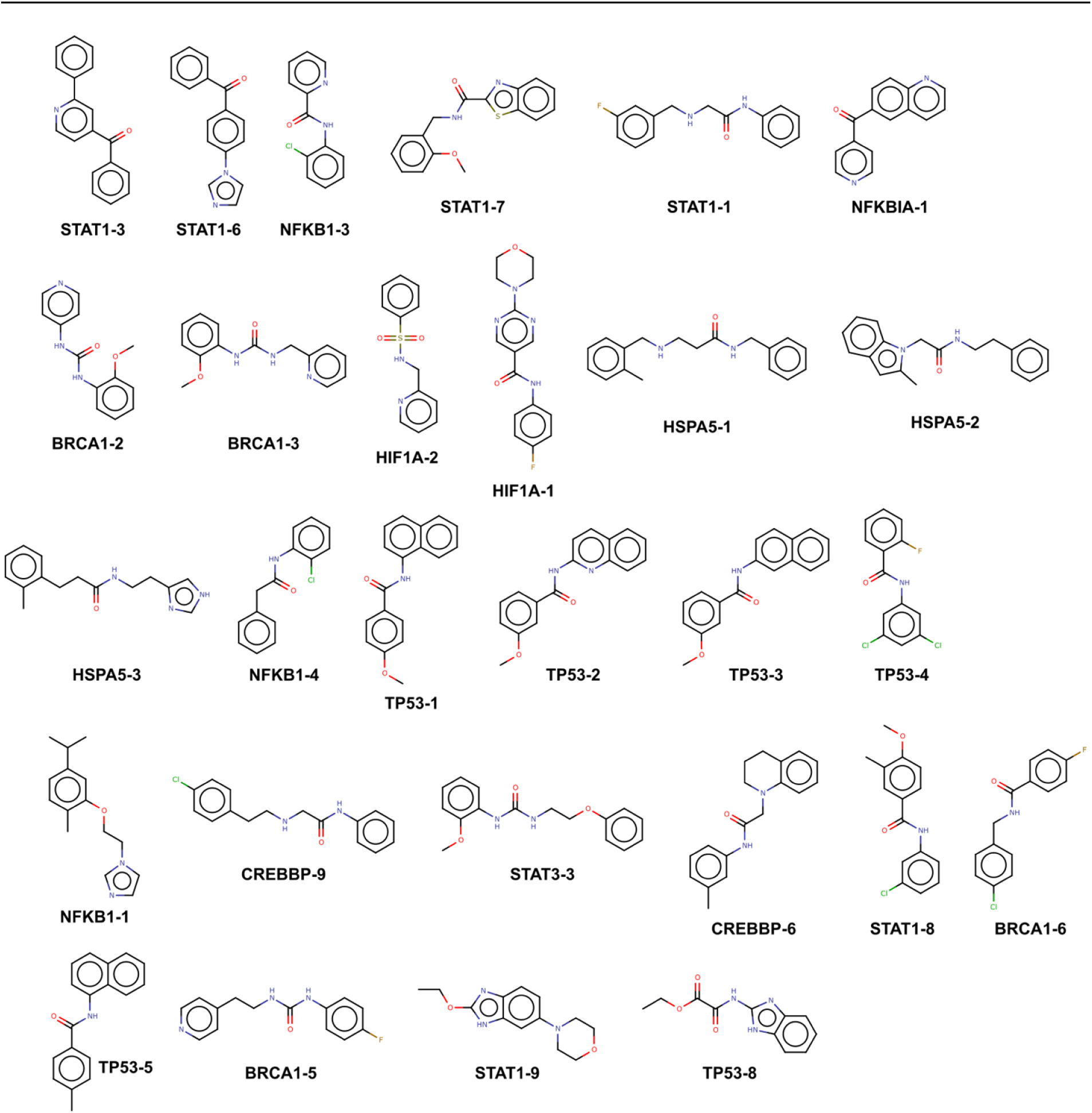
Representative chemical structures of generated compounds already in the public domain. This figure shows that compounds generated by the model used in this study are diverse and show different molecular scaffolds.

**Figure S2.**
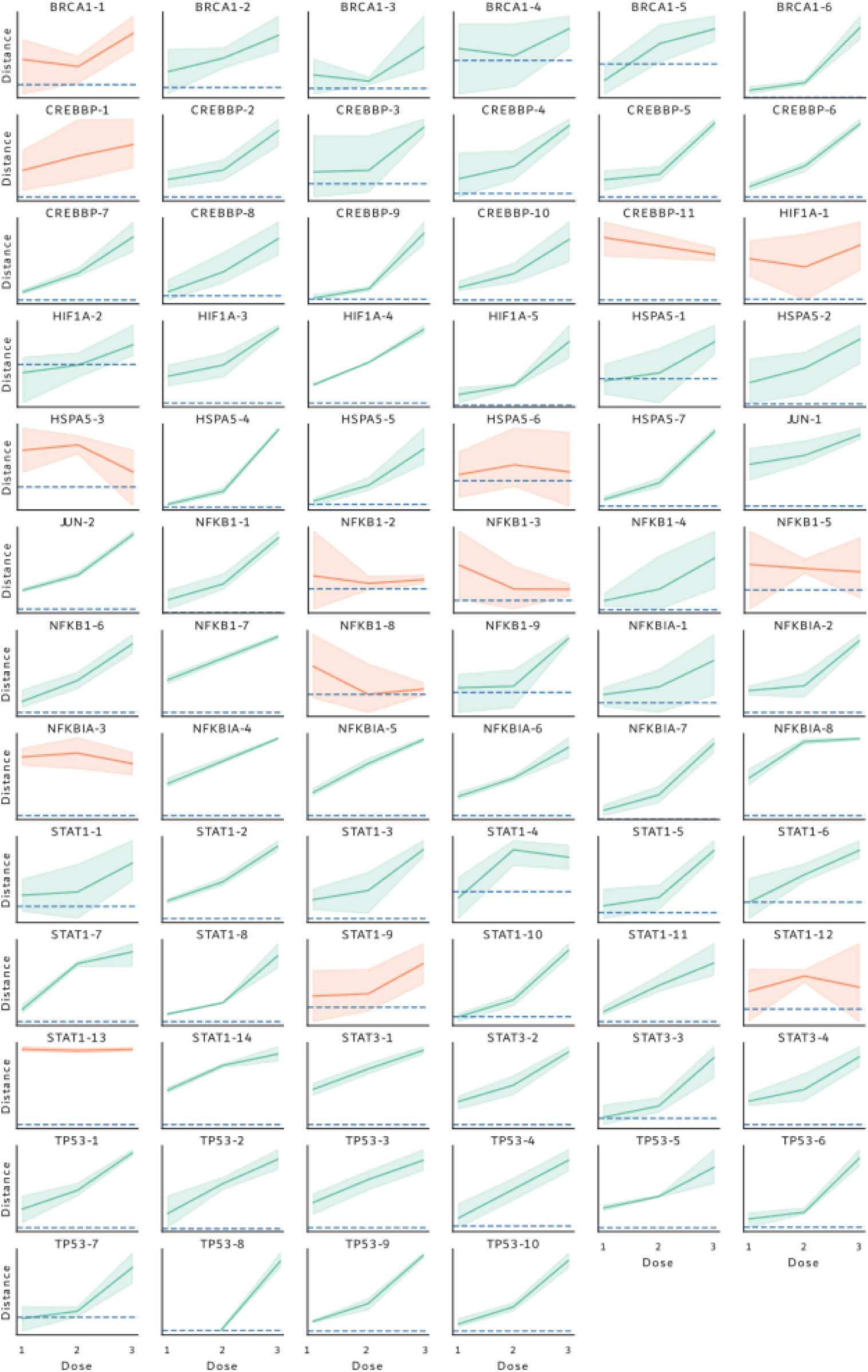
Assessment of bioactivity of de novo compounds, related to Fig. 1. A) Graphs illustrating the Mahalanobis distance to DMSO of the cell painting profiles of de novo compounds across doses. The Kendall tau test was used for classifying active compounds (depicted in green) from inactive ones (shown in orange)

**Table S1.** Number of de novo compounds synthesised per target and tested in biological assays.

| Target gene | no. of parent profiles | no. of de novo compounds | No. of compounds tested in Cell Painting | no. of compounds tested in transcriptomics | no. of compounds tested in cellular activity assay |
| --- | --- | --- | --- | --- | --- |
| BRCA1 | 2 | 6 | 6 | 6 | - |
| CREBP | 4 | 11 | 11 | 10 | - |
| HIF1A | 4 | 5 | 5 | 5 | - |
| HSPA5 | 2 | 7 | 7 | 7 | - |
| JUN | 2 | 2 | 2 | 2 | - |
| NFKB1 | 3 | 9 | 9 | 9 | - |
| NFKBIA | 4 | 8 | 8 | 6 | - |
| STAT1 | 5 | 14 | 14 | 14 | - |
| STAT3 | 3 | 4 | 4 | 4 | - |
| TP53 | 2 | 10 | 10 | 10 | 10 |

**Table S2.**
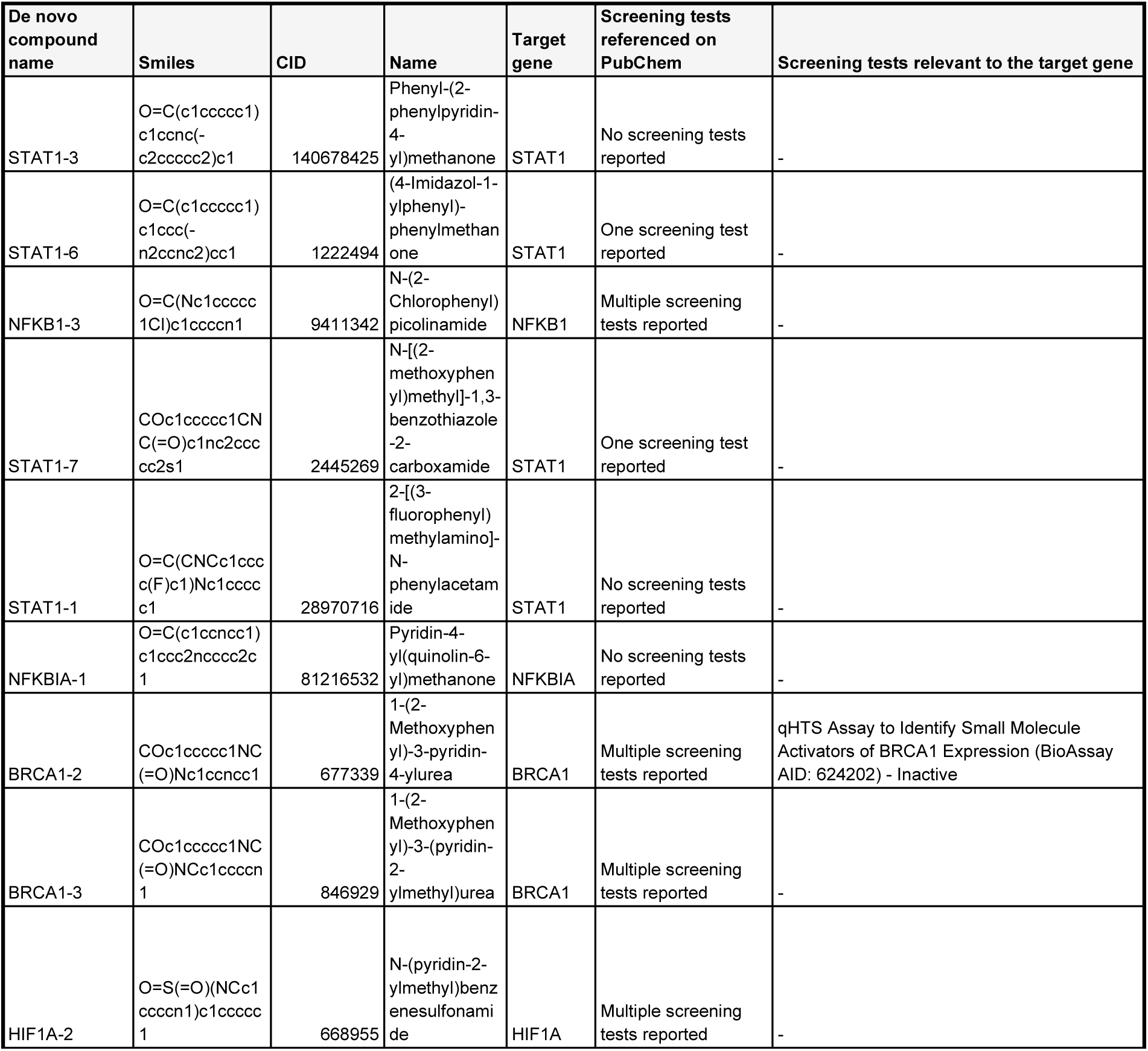
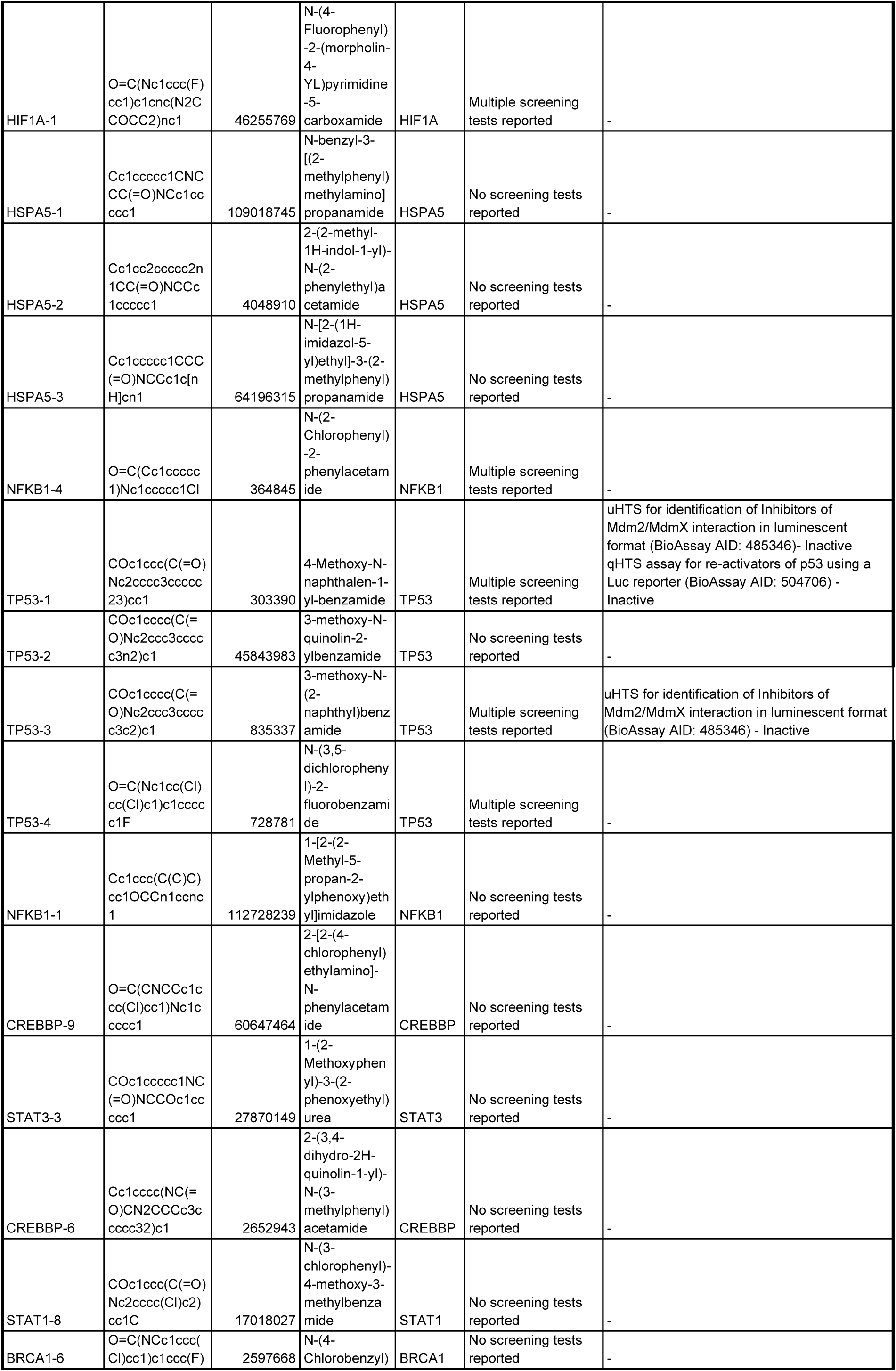
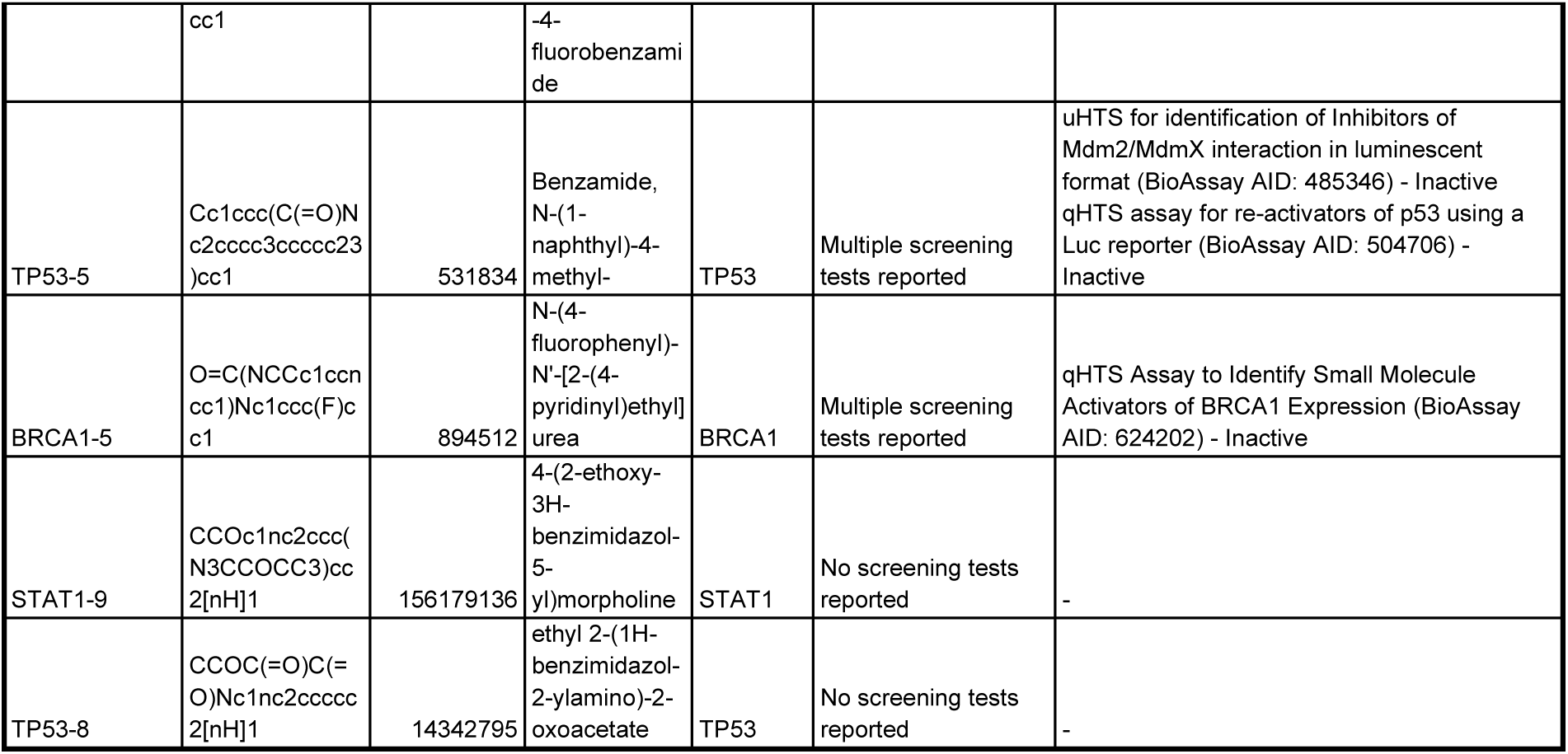
De novo compounds with existing records in PubChem with information on high-throughput biological assays annotated in the PubChem database.

## Notes

### Competing Interest Statement

The authors have declared no competing interest.

